# Energy metabolism adaptations and gene expression reprogramming in a cellular MAFLD model

**DOI:** 10.1101/2021.11.08.467719

**Authors:** Tianran Zhou, Cagla Cömert, Xiaoyu Zhou, Lin Lin, Lars Bolund, Johan Palmfeldt, Guangdong Tong, Yonglun Luo, Peter Bross

## Abstract

Mitochondrial dysfunction plays a critical role in metabolic associated fatty liver disease (MAFLD). This study aims to characterize mitochondrial dysfunctions in a human MAFLD Huh7 cell model triggered by free fatty acid (FFA) (palmitate and oleate) overload for 24 hours. We investigate its impact on cellular energy metabolism and identify potential targets for MAFLD treatment. FFA-treated cells displayed an accumulation of lipid droplets and slightly decreased viability but no significant changes in mitochondrial superoxide levels. Bioenergetic analysis showed a shift to more respiration and less glycolytic fermentation. Comprehensive transcriptomics and proteomics analyses identified changes in the expression of genes prominently involved in fatty acid handling and metabolism. The expressions of seven genes were consistently and significantly (*p* < 0.05) altered (4 upregulated and 3 downregulated genes) in both proteomics and transcriptomics. The FFA-treated Huh7 cell model is an appropriate *in vitro* model to study fatty acid metabolism and suitable to investigate the role of mitochondria, glycolysis, and multiple metabolic pathways in MAFLD. Our comprehensive analyses form a basis for drug discovery and screening using this model.

## INTRODUCTION

Metabolic (dysfunction) associated fatty liver disease (MAFLD) (Eslam *et al*, 2020), previously called non-alcoholic fatty liver disease (NAFLD), comprises a spectrum of chronic liver diseases characterized by excessive cytoplasmic retention of triglycerides (Anstee *et al*, 2019). The deposition of free fatty acids (FFA) in liver leads to simple steatosis of hepatocytes, but the underlying mechanisms are still unclear (Li *et al*, 2017). If left untreated, simple steatosis may progress to liver fibrosis and even hepatocellular carcinoma (Santhekadur *et al*, 2018). MAFLD affects the quality of life of patients, and places a heavy burden on the family, the healthcare system and the economy (Younossi, 2019). The pathogenesis of MAFLD is still largely unclear, and this is the major reason why there are currently still no FDA recommend drugs for MAFLD treatment (Eslam *et al*, 2020). Weight loss and bariatric surgery are the treatment options for MAFLD, but both have inevitable limitations (Younossi *et al*, 2018).

Mitochondrial metabolic dysfunction has been associated with MAFLD, and it has been indicated that it may play a key role in the pathogenesis of MAFLD (Mansouri *et al*, 2018). Hepatic changes in fatty acid β-oxidation, mitochondrial oxidative phosphorylation (OXPHOS), and reactive oxygen species (ROS) production are considered to be central manifestations of mitochondrial dysfunctions triggering the progression of MAFLD (Gariani *et al*, 2017; Chen *et al*, 2020; Piccinin *et al*, 2019). The disease proceeds from simple fatty liver disease, via steatohepatitis to fibrosis, and can finally develop into hepatocellular carcinoma (HCC) (Cholankeril *et al*, 2016; Hardy *et al*, 2016). Cell and animal models mimicking different stages of the disease allow studying phenotypic changes and pathogenetic mechanisms of MAFLD. So far, the mechanisms that link hepatic fatty acid accumulation to disturbances in the expression of nuclear and mitochondrial DNA encoded proteins and in turn, lead to mitochondrial dysfunction are sparsely elucidated.

In order to characterize the mitochondrial dysfunction phenotypes and mechanisms in FFA-induced MAFLD, we used Huh7 cells treated with a high dose of FFA consisting of the long-chain saturated fatty acid palmitate (C16:0) and the monounsaturated fatty acid oleate (C18:1) as an *in vitro* model. Fatty acids (molar ratio of palmitic acid : oleic acid = 1:2) were conjugated to albumin, and albumin was used as a vehicle to provide FFAs for cellular uptake. A previous study demonstrated responses induced by this FFA treatment ranging from increased levels of inflammation markers, increased levels of cellular hydrogen peroxide, increased apoptosis tendency, and the production of fibrogenic cytokines (Chavez-Tapia *et al*, 2012). We here further characterize the FFA effects by performing real-time cell metabolic analysis, high-throughput imaging flow cytometry (HTIFC), as well as global proteomics and transcriptomics analyses. We discuss our novel findings in relation to the literature. We conclude that the FFA treated Huh7 cell model is an appropriate *in vitro* model to study the role of mitochondria, fatty acid metabolism, glycolysis, and multiple metabolic pathways in MAFLD. Our molecular and phenotypic characterization can form the basis for using it in drug screening.

## MATERIALS AND METHODS

### Cell Culture and Treatment

Human hepatocellular carcinoma cell line (Huh7) (Cheng *et al*, 1993) was cultured in Dulbecco’s modified eagle medium (DMEM) with high glucose (Sigma) supplemented with 10% fetal bovine serum (FBS) (Biowest), 2 mM L-Glutamine (Sigma), 10,000 U/mL penicillin and 10 mg/mL streptomycin (Sigma) at 37 °C under 5% CO_2_, in a 95% humidified atmosphere. Cells were exposed for 24 h to 1200μM exogenous fatty acid mixture containing palmitic (P0500, Sigma) and oleic acid (O1008, Sigma). The mixture was freshly prepared before each experiment. In short, fatty acid stock solution was added to prewarmed complete medium (molar ratio of palmitic acid : oleic acid = 1:2, complexed with 300μM fatty acid-free bovine serum albumin (BSA) (A8806, Sigma) to induce MAFLD at the stage of steatohepatitis as previously described (Chavez-Tapia *et al*, 2012).

### Cell Viability and *in Vitro* Cytotoxicity Assay

Cell viability was assessed using the CCK-8 kit (Sigma) for quantitation of viable cell number in proliferation and cytotoxicity assays. Briefly, 8 × 10^3^ cells/well were seeded in a 96-well microplate and pre-incubated for 24 h. The medium was then removed, and cells were treated with medium with or without FFA for another 24 h. Cells were washed twice with DMEM medium followed by the addition of 10 μL of CCK-8 in 100μL DMEM medium and incubation at 37 °C for 80 min. The absorbance at 450 nm was measured by Synergy H1 Hybrid Multi-Mode Reader (BioTek) and Gen5 software.

### Oil Red O and Coomassie Brilliant Blue Staining

Cells were washed twice in PBS and fixed using 10% formalin for 1 h. Cells were then incubated in 60% isopropanol for 5 min. Isopropanol was discarded, and cells were incubated in Oil Red O Working Solution (Sigma) for 20 min. Oil Red O Working Solution was prepared by mixing 0.5% Oil Red O in isopropanol and water in the ratio of 3:2. After discarding the Oil Red O solution, the cells were washed 5 times with sterile ultrapure water. The absorbance at 510 nm was measured by Synergy H1 Hybrid Multi-Mode Reader (BioTek) and Gen5 software. Cells were counterstained with 0.25% Coomassie Brilliant Blue R-250 dye (AKH Reagent) for 5 min and washed 3 times with sterile ultrapure water to visualize the cytoskeleton. Images were taken by a light microscope camera (ZEISS), both with a 10x and a 40x objective, respectively (Fig. 1B).

**Figure 1.**
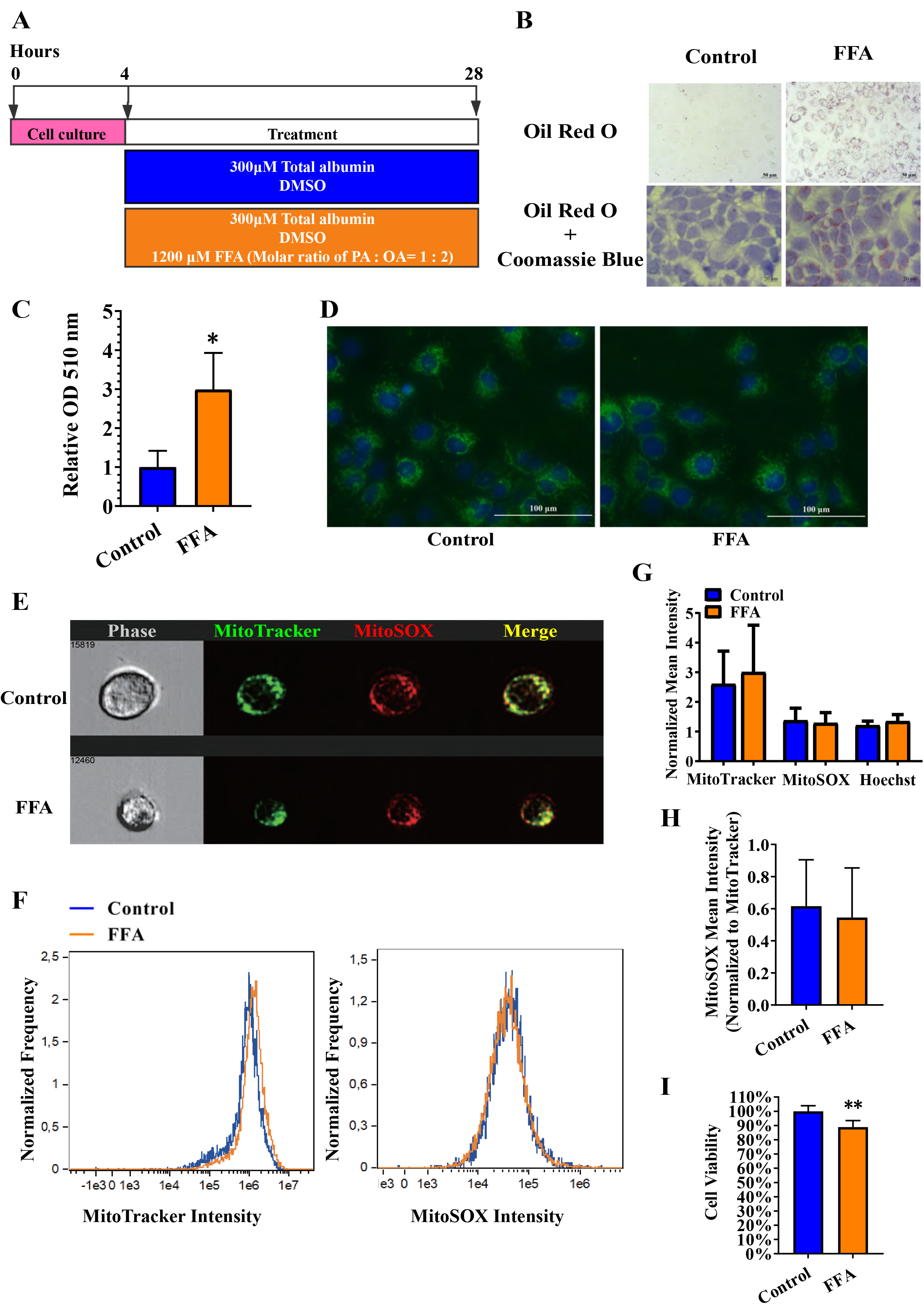
Effects of FFA treatment on Huh7 cells. (A) Workflow for the establishment of the *in vitro* MAFLD Huh7 cell model. PA: palmitic acid, OA: oleic acid. (B) Light microscopy analysis (100x and 400x magnification) of lipid droplets in Huh7 cells with or without treatment with FFA for 24 h. Cells were stained by Oil Red O solution after fixing. (C) Quantification of Oil Red O staining in MAFLD model by a plate reader. Absorbance values were recorded at 510 nm and normalized to the untreated cells. n=3. (D) Fluorescent microscopy images (400x magnification) of MitoTracker Green staining of mitochondria in live Huh7 cells treated with or without FFA for 24 h. (E) Flow cytometry and microscopy analysis. Control and model cell were stained with MitoSOX Red and MitoTracker Green as described in materials and methods and analysed using Imaging flow cytometry. The left panel shows phase-contrast images of the representative cells, and the respective fluorescence channel pictures are shown in the MitoTracker Green, MitoSOX, and merge channels. (F) Histogram of average MitoTracker Green and MitoSOX Red fluorescent intensity quantitations in Huh7 cells with or without treatment with FFA for 24 h from two representative high-throughput imaging flow cytometry (HTIFC) experiments. (G) Quantification of average fluorescence intensity of MitoTracker Green per cell, MitoSOX per cell, and Hoechst 33342 per cell in Huh7 cells with or without treatment with FFA for 24 h. n=4. (H) Quantification of average MitoSOX fluorescence intensity related to MitoTracker Green in Huh7 cells with or without treatment with FFA for 24 h. n=4. All numeric results are expressed as the mean value of triplicate samples normalized to the untreated cells ± SD (error bars). \**p* < 0.05, \*\**p* < 0.01. (I) Viability of Huh7 cells measured by Cell Counting Kit - 8 assay after a 24-h exposure to FFA. n=3.

### Seahorse analysis of cellular bioenergetics

The real-time cellular oxygen consumption rate (OCR) and extracellular acidification rate (ECAR) were measured using the Seahorse XFe96 extracellular flux analyzer (Seahorse Bioscience). Experiments were performed according to the manufacturer’s instructions with minor modifications as specified below. OCR and ECAR were measured using Seahorse XF Cell Mito Stress Test Kit (Agilent Technologies). 1 × 10^4^ cells/well were plated onto 7 μg/mL Poly-D-Lysine coated (Ruhanen *et al*, 2017) Seahorse XF cell culture microplates and allowed to attach for 4 h. The medium was then removed, and cells were treated with DMEM medium with FFA or control medium for 24 h. Based on pilot experiments, we used the following procedure: after baseline measurements, oligomycin (1.5 μM), FCCP (0.65μM), and rotenone plus antimycin A (Rot/AA) (0.5μM) were sequentially injected into each well at the indicated time points. Three consecutive measurements were performed for each step/injection. After OCR and ECAR measurement, cells were stained by 2.5 μM Hoechst 33342 for 55 min. Hoechst fluorescence was recorded by Cytation 1, cell counts per well were calculated with Seahorse XF Imaging and Cell Counting Software, and OCR and ECAR data were normalized by cell number. Data were further analyzed by Seahorse XFe Wave software and exported to Microsoft Excel. OCR is reported in pmols/min/1000 cells and ECAR in mpH/min/1000 cells.

MitoATP Production Rate (pmol ATP/min/1000 cells), glycoATP Production Rate (pmol ATP/min/1000 cells), total ATP Production Rate (pmol ATP/min/1000 cells), % Glycolysis, % Oxidative Phosphorylation, XF ATP Rate Index were calculated from normalized real-time OCR (pmols/min/1000 cells) and ECAR (mpH/min/1000 cells) data from Seahorse XF Cell Mito Stress Test according to 11 equations in manufacturer’s ATP Rate Assay instructions.

Glycolytic rate assay parameter, including basal glycolysis (glycoPER) (pmol/min/1000 cells), basal proton efflux rate (PER) (pmol/min/1000 cells), % PER from glycolysis (basal) (%), and basal mitoOCR/glycoPER data were calculated from real-time OCR, ECAR and predetermined CO_2_ contribution factor according to manufacturer’s Glycolytic Rate Assay user guide.

### MitoTracker Green Live Cell Staining

Huh7 cells were incubated at 37 °C with prewarmed (37 °C) 200 nM MitoTracker Green (ThermoFisher) and 2.5μM Hoechst 33342 (ThermoFisher) in Hank’s balanced salt solution (HBSS) with calcium and magnesium (Gibco) simultaneously for 45 min. The staining solution was removed, and cells were added HBSS. Live cell images were recorded immediately by a fluorescent microscope (EVOS® FL) with a 40x objective (Fig. 1D).

### Live Cell Staining and High-throughput Imaging Flow Cytometry (HTIFC)

Firstly, live cells were washed twice with warm PBS, and stained with prewarmed 5 μM MitoSOX Red (ThermoFisher) for 10 min at 37 °C. Secondly, cells were then washed twice, and stained with prewarmed 0.5% BSA/PBS containing 100 μM MitoTracker Green (ThermoFisher) and 3.3 μM Hoechst 33342 (ThermoFisher) and incubated for 45 min at 37 °C. Thirdly, cells were washed, lifted with trypsin/EDTA, centrifuged, resuspended with 0.5% BSA/PBS. Compensation controls were prepared for each fluorochrome to make a standard compensation matrix file. A minimum of 10,000 Huh7 cells were acquired for analysis.

Samples were acquired using an ImageStreamX Mark II multispectral imaging flow cytometer (Amnis) equipped with INSPIRE ImageStreamX System software. Data were analyzed using IDEAS image data exploration and analysis software version 6.3 (Amnis).

### RNA Library Construction and mRNA-sequencing

Total RNA from control (n=3) and model (n=3) Huh7 cells were isolated by TRIZOL reagent (Thermo Scientific), and the RNA concentration, purity and integrity were assessed by the Synergy H1 Hybrid Multi-Mode Reader (BioTek) and 2100 Bioanalyzer (Agilent Technologies). Total RNA samples were stored at −80 °C until further analysis.

The construction and deep sequencing of RNA libraries were accomplished with the assistance of BGI Europe Co., Ltd. Briefly, by oligo(dT) selection, mRNA was enriched and purified from total RNA, followed by RNA fragmentation, reverse transcription, end repair, add A, and adaptor ligation. After PCR amplification, the double-stranded PCR products were subjected to heat separation, cyclization (circularized by the splint oligo sequence), DNA nano ball synthesis. The single-stranded circular DNAs were formatted as the final library for library quality assessment and subsequent PE100 (paired-end 100) sequencing. The mRNA-sequencing was performed on a well-established DNBSEQ/MGISEQ2000 Technology Platform (Rao *et al*, 2020; Yue *et al*, 2020; Patterson *et al*, 2019) (BGI, Copenhagen, Denmark).

### Transcriptome Pipeline for mRNA-seq and Gene-Level Differential Expression Analysis

The upstream pipeline includes quality control, adapter trimming, quality filtering, genome alignment, reads deduplication, and estimating gene expression levels.

After removing duplicate reads from bam files, counting reads were used as a measure of gene expression by featureCounts, raw data counts were normalized separately by transcripts per million (TPM) (reference of R script: https://github.com/t-arae/ngscmdr/blob/master/R/calc_rpkm.R), DESeq2 (version 1.30.1), and EdgeR (version 3.32.0) with trimmed mean of M value (TMM). Raw count data from RNA-seq were normalized by TPM (Wagner *et al*, 2012) method for gene expression heatmap visualization. TMM (Robinson & Oshlack, 2010) was calculated by EdgeR for further gene set enrichment analysis (GSEA). HISAT2 software (version 2.2.1) was used for mapping mRNA-sequencing reads. Deduplication (Dedup) was performed by the Picard tool (version 2.23.8). GENCODE V35 was the selected annotation gene sets, and GRCh38 primary assembly version was the selected human genome. The number of annotated genes in the selected gene sets (GENCODE V35) was 60,715 (including 19,983 coding genes, 16,899 non-coding genes, and 1,881 pseudogenes). Gene expression levels were quantified with the above reads/annotations by featureCounts without gene types filtering. During the differential expression analysis, the gene types and symbols were annotated and added in the raw counts data, then coding genes were selected for further analysis while TPM values of all genes were used for GSEA analysis.

Raw count data normalization, protein-coding genes selection, and differential expression analysis were performed by DESeq2 R/Bioconductor package (Anders & Huber, 2010). Fold change, log_2_(fold change) and *p*-value were calculated, gene type and other annotations were reported by biomaRt. *P* < 0.05 were selected as the cutoff criteria to identify differentially expressed genes for the DESeq2 method.

### Gene Ontology (GO) Analysis and GSEA Analysis

Differentially expressed gene analysis was performed using DAVID bioinformatics resources (Huang *et al*, 2009a, 2009b) using all *Homo sapiens* genes as background. Categories within the Uniprot keywords, GO (BP, CC, and MF) KEGG were extracted and replotted. Count ≥ 2, EASE score ≤ 0.1 were used as thresholds for DAVID functional annotation analysis.

The long list of GO BP terms was summarized and visualized by REVIGO (http://revigo.irb.hr/) (Supek *et al*, 2011). Redundant GO BP terms were removed, and the remaining terms were visualized in the semantic similarity-based interactive graph by REVIGO. The interactive graph was exported to an XGMML file and further processed by the Cytoscape 3.8.2 program (Su *et al*, 2014; Shannon *et al*, 2003).

Count data normalized by the TMM method were used for the GSEA expression data set. GSEA was performed using the GSEA 4.1.0 software (Mootha *et al*, 2003; Subramanian *et al*, 2005) (https://www.broadinstitute.org/gsea/) with permutation type = geneset, enrichment statistic = weighted, metric for ranking genes = signal to noise, #permutation = 1000. Several gene set collections in the Molecular Signatures Database (MSigDB) (Liberzon *et al*, 2011; Subramanian *et al*, 2005) were used for GSEA, including hallmark gene sets (Liberzon *et al*, 2015), curated gene sets, and ontology gene sets.

### Sample Preparation for Proteomics

Protein samples were prepared for labelling by TMT sixplex isobaric mass tagging kit (Thermo Scientific) according to the manufacturer’s protocol. In short, 80 μg of total protein was reduced and alkylated with tris(2-carboxyethyl)phosphine and iodoacetamide, followed by overnight acetone precipitation. After redissolving the pellet, proteins were digested overnight with trypsin (Promega). The resulting peptides were labelled with the TMT sixplex isobaric mass tag, and the six samples – three treated and three control samples– were pooled. Further treatment and fractionation were performed as previously described (Palmfeldt *et al*, 2009). The pool of labelled peptides was dissolved in 10 mM phosphoric acid, pH 3.0 and 25% acetonitrile before strong cation exchange (SCX) purification using a SCX cartridge (Phenomenex). After washing, drying, elution procedure, and drying by a SpeedVac vacuum concentrator overnight, vacuum dried labelled peptide pools were rehydrated and fractioned into ten fractions by isoelectric focusing with an immobiline drystrip pH 3-10, 18 cm (GE Healthcare). Peptides were extracted and dissolved in 5% acetonitrile, 0.5% trifluoroacetic acid, and further purified by C18 spin columns (Thermo Scientific) according to the manufacturer’s protocol. After elution, samples were dried by a SpeedVac vacuum concentrator and stored at −20°C until nanoLC-MS/MS analyses.

### NanoLC-MS/MS and Proteomics Database Searches

Liquid chromatography-tandem mass spectrometry (LC-MS/MS) was performed, as previously described (Paternoster *et al*, 2019), on an EASY nanoLC-1000 coupled to Q Exactive™ HF-X Quadrupole-Orbitrap™ Mass Spectrometer (Thermo Scientific). Pre-column (Acclaim PepMap 100, 75μm x 2 cm, Nanoviper, Thermo Scientific) and analytical column (EASY-Spray column, PepMap RSLC C18, 2μm, 100 Å, 75 μM x 25 cm) were used to trap and separate peptides using a 170 mins gradient (5-40 % acetonitrile, 0.1 % Formic acid). The MS was operated in positive mode, and higher-energy collisional dissociation (HCD) with collision energy (NCE) of 35 was applied for peptide fragmentation. Full scan (MS1) resolution was 60,000, and AGC target set at 1×10^6^ with scan range between 392-1,500 m/z. Data-dependent analysis (DDA) was applied to fragment up to 12 of the most intense peaks in MS1. Resolution for fragment scans (MS2) was set at 45,000 with the first fixed mass at 110 m/z and AGC target at 1×10^5^. Dynamic exclusion was set at 15 seconds, and unassigned and +1 charge states were excluded from fragmentation. Each sample was LC-MS/MS analyzed twice. Peptides identified with more than nine peptide spectrum matches (PSM) in the first analysis were excluded from fragmentation in the second analysis. All 20 (=2×10) LC-MS analyses of the study were merged and submitted for database search for protein identification and quantification in Proteome Discoverer 2.3 (Thermo Scientific) using the Sequest algorithm, with 20,422 reviewed *Homo sapiens* Uniprot sequences as reference proteome. Precursor and fragment mass tolerance were set at 10 ppm and 20 mmu, respectively. Oxidation of methionine was set as dynamic modification, and static modifications were carbamidomethylation of cysteines and TMT sixplex labels on lysine and peptide N-terminus. The co-isolation threshold was set at 45 %, and the identification false discovery rate was set to 0.01.

### Bioinformatics Analysis of Differential Gene Expression at protein level

*P* < 0.05 and |log_2_Fold change| > 0.263034 were selected as the cutoff criteria to identify differential expression proteins (Levin, 2011).

Donut charts were drawn, and differentially expressed proteins with statistical significance were highlighted in the volcano plots using GraphPad Prism 9 software. Hierarchical clustering was performed using Morpheus (https://software.broadinstitute.org/morpheus). Venn Diagrams were performed using InteractiVenn (Heberle *et al*, 2015).

Uniprot Keywords, GO analysis, including biological process (BP), cellular component (CC) and molecular function (MF) groups, and Kyoto encyclopedia of genes and genomes (KEGG) pathways enrichment analysis of differentially expressed proteins were performed using DAVID bioinformatics resources (Huang *et al*, 2009a, 2009b) using all detected high-confidence proteins as background. Count ≥ 2, a modified Fisher Exact *p* (EASE score) ≤ 0.1 were used as thresholds for DAVID functional annotation analysis.

MitoCarta3.0 datasets, which contains detailed information on the 1136 mitochondrial human genes, including sub-mitochondrial localization and pathways (Rath *et al*, 2021), were used to identify human mitochondrial proteins.

The long lists of GO BP terms were summarized and visualized by REVIGO (http://revigo.irb.hr/) (Supek *et al*, 2011). Redundant GO BP terms were removed, and the remaining terms were visualized in the semantic similarity-based interactive graph by REVIGO. The interactive graph was exported to an XGMML file and further processed by the stand-alone Cytoscape 3.8.2 software (Su *et al*, 2014; Shannon *et al*, 2003).

Pivot chart was used to show the frequency distribution of log_2_ (fold change) for transcriptomics and proteomics. TPM ≥ 1 was used as a threshold for fold change for the transcriptomics dataset only for genes quantitated commonly in transcriptomics and proteomics analyses. The pivot table was generated using Microsoft Excel, and pivot graph was generated using GraphPad Prism 9.

Protein-protein interaction (PPI) clustered network of the 8 co-regulated genes and proteins was generated and exported by STRING 11.0 online tool (Szklarczyk *et al*, 2019; von Mering *et al*, 2003) (https://string-db.org/). Network type = full STRING network, required score = low confidence (0.150), FDR stringency = medium (5%), organism: *Homo sapiens*, network clustering *=* kmeans clustering, number of clusters = 3 were used as thresholds and parameters for STRING analysis.

### Statistical analysis

Normal distribution data were expressed as means ± SD around the mean, non-normal distribution data were expressed as percentiles. It was checked whether the data conformed to the normal distribution. For data conforming to the normal distribution, non-pairwise comparisons were performed using the Student’s *t*-test. A non-parametric test was used for the non-normally distributed data. Calculations were performed using SPSS 20.0. A *p* of < 0.05 was considered statistically significant. Hierarchical clustering analysis was performed using Morpheus (https://software.broadinstitute.org/morpheus). Correlation analysis was performed using GraphPad Prism 9.

## RESULTS

### Establishment of the Huh7 cell MAFLD model

To investigate the cellular response to FFA treatment, we established a cell-based MAFLD model that has previously been described by Chavez-Tapia *et al* (Chavez-Tapia *et al*, 2012). For the model, the human liver cancer cell line Huh7 was treated for 24 hours with a 1:2 mixture of palmitic and oleic acid (total FA concentration 1200 μM) bound to BSA (Fig. 1A). Oil red O staining clearly showed an increased accumulation of lipid droplets within FFA-treated Huh7 cells (Fig.1B). Quantitation of the Oil red O staining showed a significant increase (199%) of lipid staining in FFA-treated cells (Fig. 1C). Taken together, this establishes that the FFA treatment regimen generates a cellular MAFLD model.

To get an overview of the effects of FFA treatment on mitochondria, we first performed fluorescence microscopy imaging of mitochondria with MitoTracker Green. This indicated no gross changes in mitochondrial size and morphology in FFA-treated cells (Fig. 1 D). Then, to study mitochondrial dysfunction-related phenotypes, we performed high-throughput single-cell imaging using MitoTracker Green staining for mitochondria volume, MitoSOX Red staining for mitochondrial superoxide, and Hoechst 33342 staining for cell apoptosis. Our results showed that FFA-treatment had no significant effect on the mitochondrial volume and caused no significant changes in superoxide levels, both as mean intensity and mean intensity normalized to MitoTracker Green staining (Fig. 1 G, H). Hoechst 33342 staining, an indicator of apoptosis (Crowley *et al*, 2016; Zhu *et al*, 2020; Reynolds *et al*, 1996), was also not significantly increased in MAFLD model cells (Fig. 1 G). However, CCK8 analysis indicated a slight (11%) but statistically significant decrease in viability of the FFA-treated cells (Figure 1 I). This suggests that there are no effects on overall mitochondrial parameters but that cell viability is impaired.

### Effects of FFA treatment on cellular bioenergetics in the Huh7 cell MAFLD model

Firstly, we wondered whether the MAFLD model cells displayed changes in key parameters of mitochondrial and cellular energy metabolism. To this end, we measured the OCR and proton excretion rate (PER) using the Seahorse XFe96 metabolic analyzer and the MitoStress Test protocol (see materials and methods). Our data showed that FFA-treated cells displayed increased oxygen consumption rates (Fig. 2A) and decreased proton excretion rates (Fig. 2A). Calculation of key parameters revealed that FFA treatment induced a significant increase (*p* = 0.047) in basal respiration (Fig. 2B) while there was no significant increase in maximal respiration (*p* = 0.391) (Fig. 2B). We observed no significant differences in non-mitochondrial oxygen consumption and proton leakage in the MAFLD model (data not shown).

**Fig. 2.**
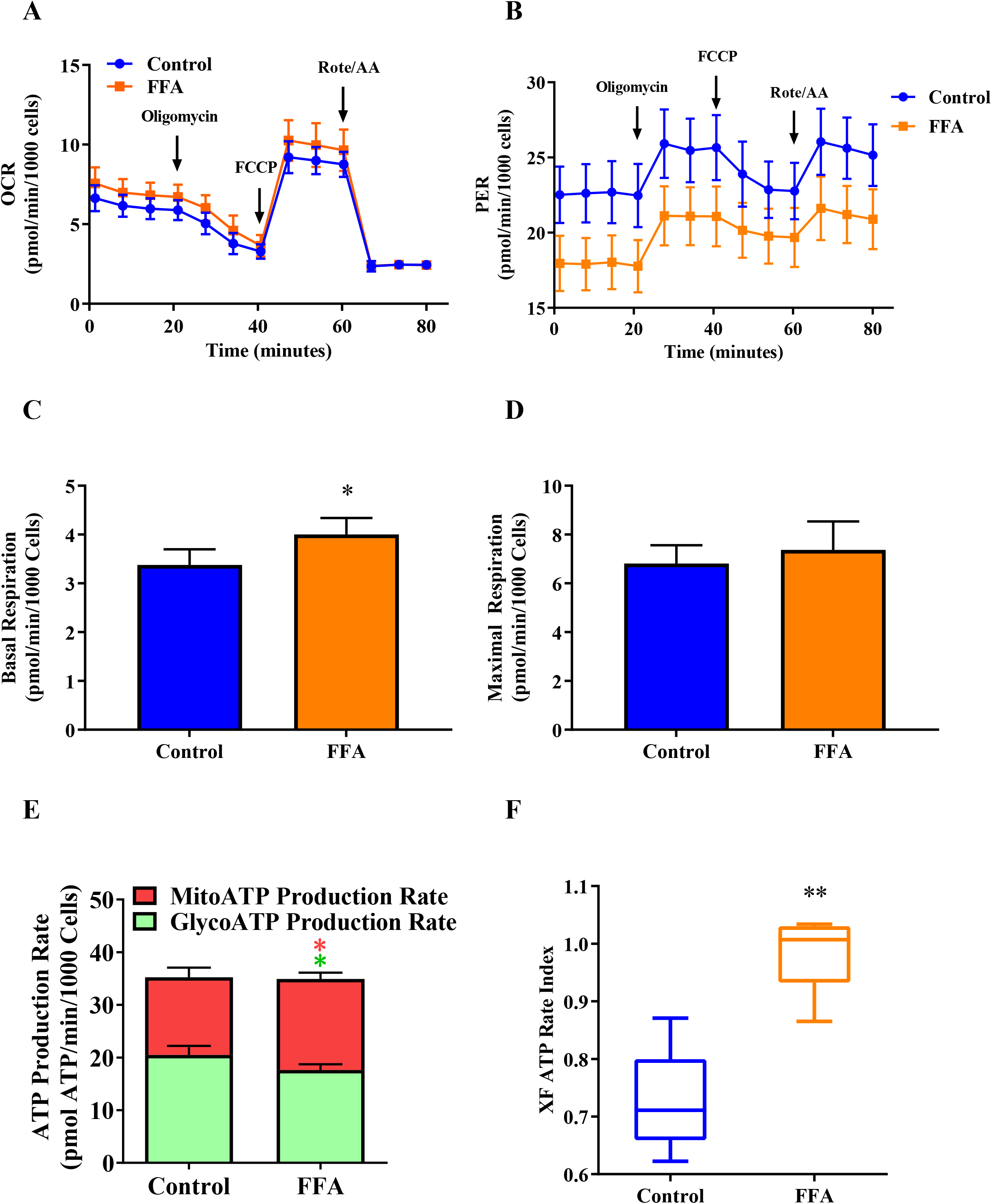
Key parameters of cellular energy metabolism in FFA-treated Huh7 cells and controls. Real time oxygen consumption rates (OCR) and proton excretion rates (PER) were measured in live Huh cells using a Seahorse XFe96 metabolic analyzer. (A) and (B) Representative OCR and PER curves, respectively, for Huh7 cells with or without FFA treatment. Using the Cell Mito Stress Test assay, Oligomycin, Carbonyl cyanide-4 (trifluoromethoxy) phenylhydrazone (FCCP), Rotenone, and Antimycin were injected at the indicated time point. Data were normalized by cell numbers determined by Hoechst staining. (C) Basal respiration. MAFLD Huh7 cells showed significantly higher basal mitochondrial respiration. (D) Maximal respiration. MAFLD Huh7 cells displayed no significant change in the maximum rate of mitochondrial respiration. (E) ATP production from mitochondrial respiration and glycolysis was calculated from the OCR and PER traces as described in materials and methods. (F) Rate index graph showing ATP production from respiration/ATP production from glycolysis for FFA-treated and controls. All numeric values are expressed as the mean value of triplicate samples ± SD (error bars) (Fig 2A-E). Values in Fig. 2F are expressed as the maximum value, minimum value, 75th percentile, 25th percentile, and median. \**p* < 0.05, \*\**p* < 0.01.

Most of the cellular ATP is produced by glycolysis in the cytosol and OXPHOS by the mitochondrial respiratory chain in the mitochondria. In the condition of fatty acid overload in Huh7 cells, we investigated whether FFA treatment affects glycolytic and mitochondrial ATP production. Based on our OCR and PER data, we calculated the ATP production rates from respiration and glycolysis, respectively (see materials and methods). This showed that, while total ATP production was unchanged, mitochondrial ATP production was increased (17.07%), and glycolytic ATP production concomitantly decreased (13.95%) in the FFA treated cells (Fig. 2C). This is also illustrated by the ATP rate index, i.e. the ratio of mitochondrial ATP production divided by glycolytic ATP production (Fig. 2D).

### Effects of FFA treatment on the Transcriptome of Huh7 Cells

To identify differentially expressed protein-coding transcripts between control and FFA-treated cells, we performed mRNA-Seq analysis (Fig. 3). The donut chart in Fig. 3A shows an overview of mRNA-Seq analysis. Using the DeSeq2 method and *p* < 0.05 as cutoff criteria, 256 transcripts were identified as differentially expressed in the FFA-treated group compared to controls (81 up- and 175 down-regulated). The heat map pattern of the 256 differentially expressed transcripts is shown in Fig. 3B. The Volcano plot (Fig. 3C) displays the distribution of all 16,001 quantitated protein-coding transcripts, and the significantly upregulated (red dots) and down-regulated (blue dots) transcripts are highlighted. Enrichment analysis of the differentially expressed transcripts using DAVID identified 122 enriched GO BP terms (EASE score < 0.1) in mRNA-seq. Figure 3D lists the top 20 BP terms with gene counts and log10 (*p-*values. For KEGG pathway analysis, several metabolism-related pathways, such as fatty acid metabolism, pyruvate metabolism, adipocytokine signaling pathway, and steroid biosynthesis showed significant *p*-values (*p* < 0.05) (Fig. 3E).

**Figure 3.**
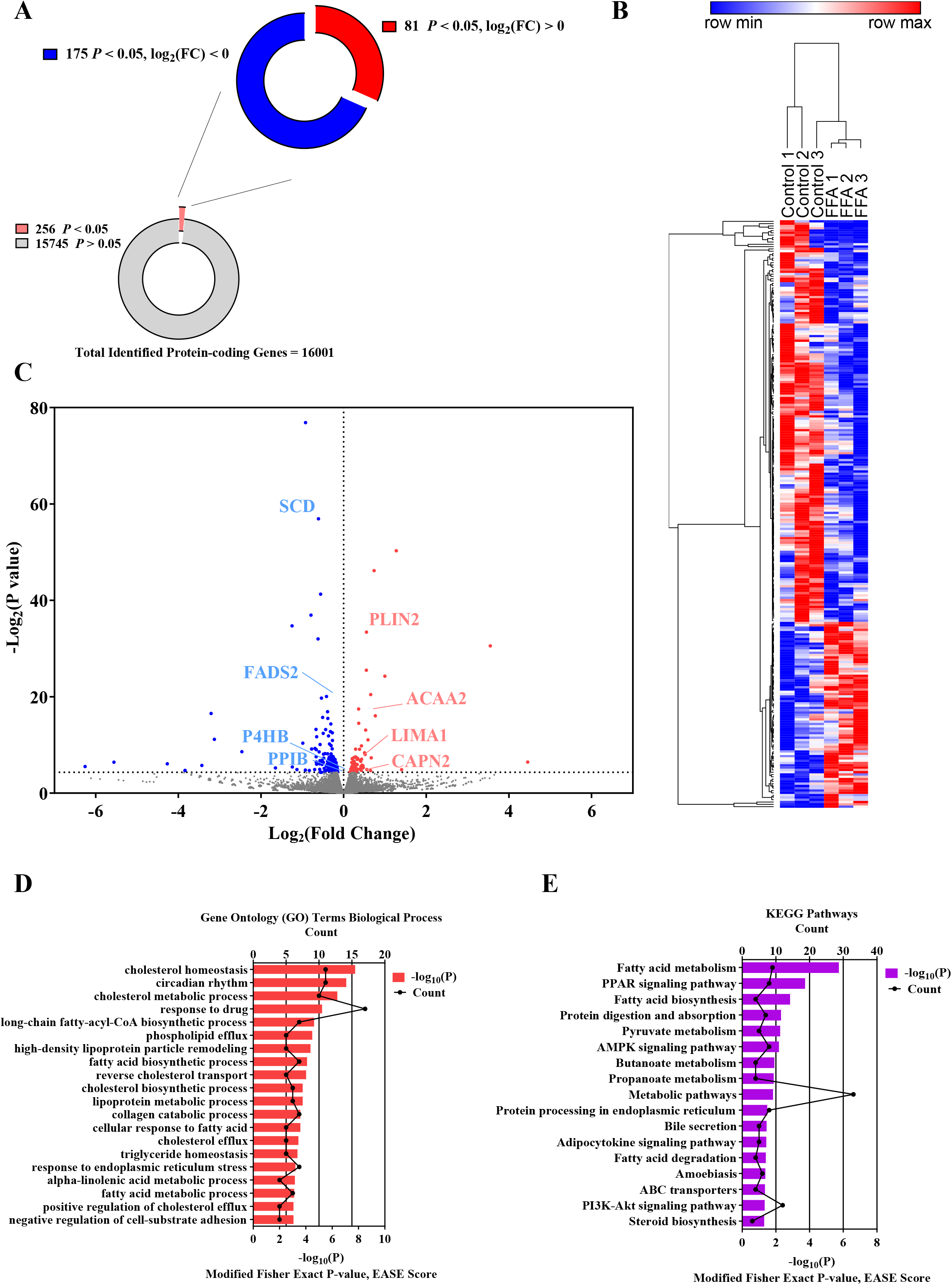
Overview of analysis of the transcriptomes FFA-treated Huh7 cells and controls. (A) Donut chart of RNA-Seq. 16,001 protein-coding genes were quantitated by RNA-Seq in control and FFA Huh7 cells, and 256 of them showed statistically significant differential expression between groups (*p* < 0.05). (B) Hierarchical clustering and heatmap of the 256 differentially expressed transcripts. (C) Volcano plot of all quantitated protein-coding transcripts. Differentially expressed genes are highlighted; red dots represent upregulated and blue dots down-regulated transcripts. Transcripts encoding proteins that were also differentially regulated at the protein level (see below) are labelled. (D-E) Bar graphs and line charts show *p*-values and count of the top 20 significantly enriched GO terms and the significant KEGG pathways (*p* < 0.05; modified Fisher’s exact test).

### Enriched Gene Sets in GSEA Analysis

Tools such as DAVID perform analysis of gene expression changes by searching for the enrichment of annotation terms in lists of genes that change in relation to a predefined threshold. This may miss important effects on pathways (Subramanian *et al*, 2005). In contrast, GSEA (Gene Set Enrichment Analysis) (Reimand *et al*, 2019; Subramanian *et al*, 2005) is a sensitive threshold-free method that analyses enrichment patterns of curated gene sets that e.g. share common biological function or regulation (Reimand *et al*, 2019).

We first used the MSigDB hallmark gene set collection that summarizes and represents specific, well-defined biological states or processes. A number of selected gene sets showed clearly enriched scores. Among these, the cholesterol homeostasis gene set was found in the top 5 upregulated hall mark gene sets in control cells, i.e., down-regulated in the model (Fig. 4A; nominal *p* = 0.0, enrichment score (ES) = −0.48). The glycolysis gene set was also one of the top 5 upregulated hallmark gene sets in control cells (Fig. 4B; nominal *p* = 0.0, ES = −0.45). This is consistent with our Seahorse analyses, which showed a downregulated glycolysis in FFA-treated cells. The chemokine receptor binding gene set was one of the significantly enriched GO MF gene sets in MAFLD model cells (Fig. 4C); nominal *p* = 0.012, ES = 0.47), and may indicate pro-inflammation in MAFLD model cells.

**Figure 4.**
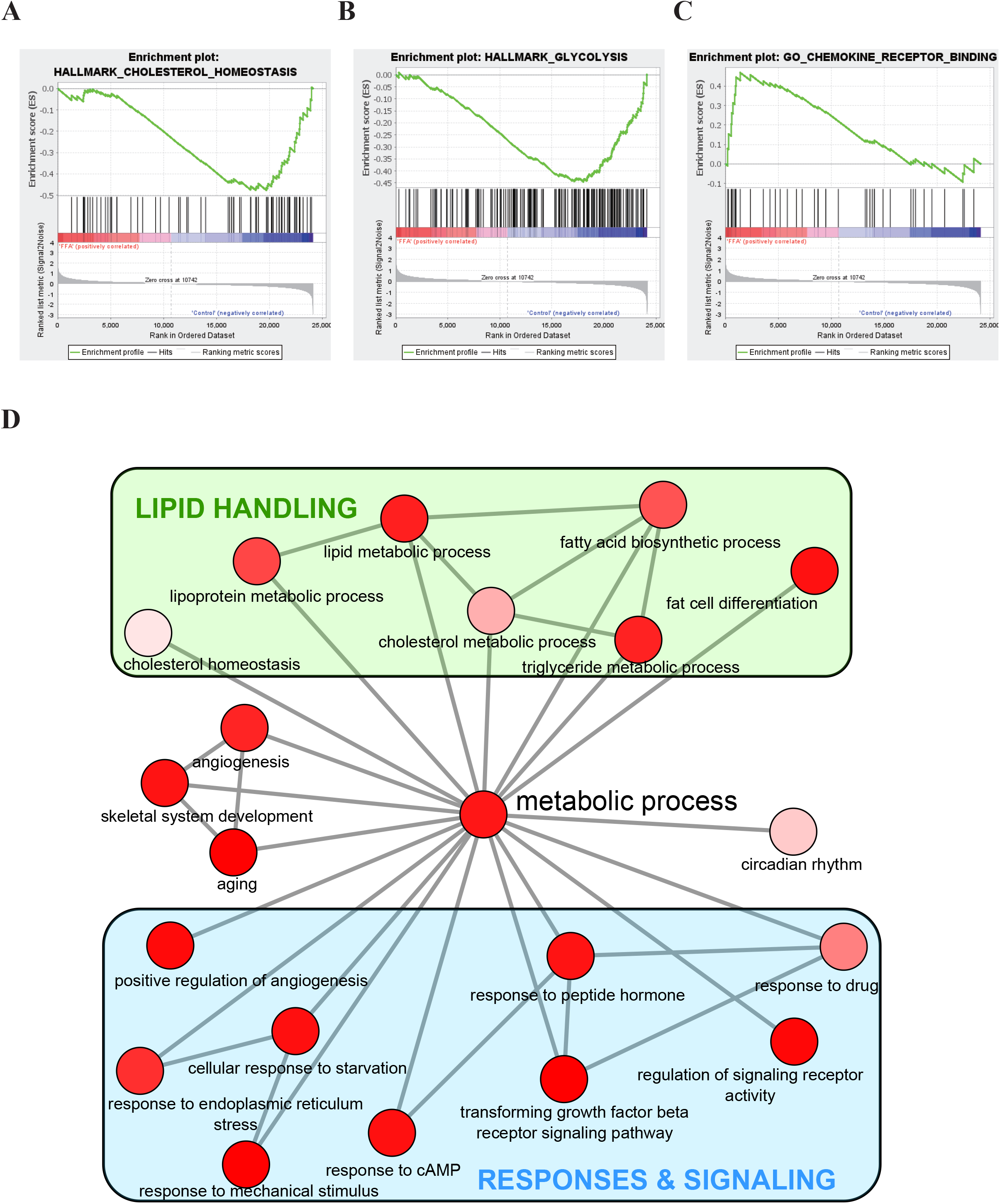
GSEA enrichment plots and redundancy-reduced GO BP terms in RNA-Seq analysis. (A-C) GSEA enrichment plots: Profile of the running ES score & positions for the cholesterol homeostasis (A), glycolysis (B), chemokine receptor binding (C) gene set members on the rank-ordered list. (D) Relationship network of redundancy-reduced enriched GO BP terms. Bubble color indicates the *p*. Highly similar GO BP terms are linked by edges in the graph. Settings: GO BP terms with EASE score (modified Fisher’s exact *p)* < 0.01 and the associated *p* were used as input for REVIGO. The REVIGO analysis was performed with the following settings: similarity = small (0.5), species = *Homo sapiens*, semantic similarity measure = Jiang and Conrath. The resulting interactive graph was edited using Cytoscape (Shannon *et al*, 2003). Consistent groups related to ‘lipid handling’ or ‘responses & signaling’ are boxed.

### Network relationships of biological process terms for the regulated genes

To get a picture of the relationships of the regulated genes in our RNA-Seq analyses, we used the REVIGO webserver to reduce redundant GO terms and visualization (Supek *et al*, 2011). By DAVID enrichment analysis, we had identified 122 enriched GO BP terms (EASE score < 0.1) in RNA-Seq. A reduced list of 51 of these terms (EASE score < 0.01) generated using REVIGO is presented as a graph in Fig. 4D. Analysis of the GO BP terms for the differentially regulated genes in RNA-Seq yielded a network with ‘Metabolic process’ linked to two larger groups of terms, one related to lipid handling and the other related to regulation and signaling (Fig. 4D).

### Effects of FFA treatment on the Huh7 cell proteome

Next, we performed proteomics analysis to identify differentially expressed proteins comparing control and FFA-treated cells and validate the transcript expression changes revealed by our mRNA-Seq analyses (Fig. 5). For the mass spectrometry-based proteomics analysis, three biological replicates of each control and FFA treated cells were cultured in parallel to the cells analyzed by mRNA-Seq. The donut chart in Fig. 5A gives an overview of the proteomics analysis. 105 proteins were found upregulated (yellow) and 67 downregulated (blue) using the cut-off criteria: |log_2_FC| > 0.263034 and *p* < 0.05). 23 of the 172 differentially expressed proteins are mitochondrial proteins annotated in MitoCarta 3.0 (Rath *et al*, 2021). Fig. 5B shows hierarchical clustering and the heatmap of the differentially expressed proteins. The volcano plot in Fig. 5C depicts fold changes and *p* of all 4110 quantitated proteins. Upregulated proteins are highlighted in red and downregulated proteins in blue.

**Figure 5.**
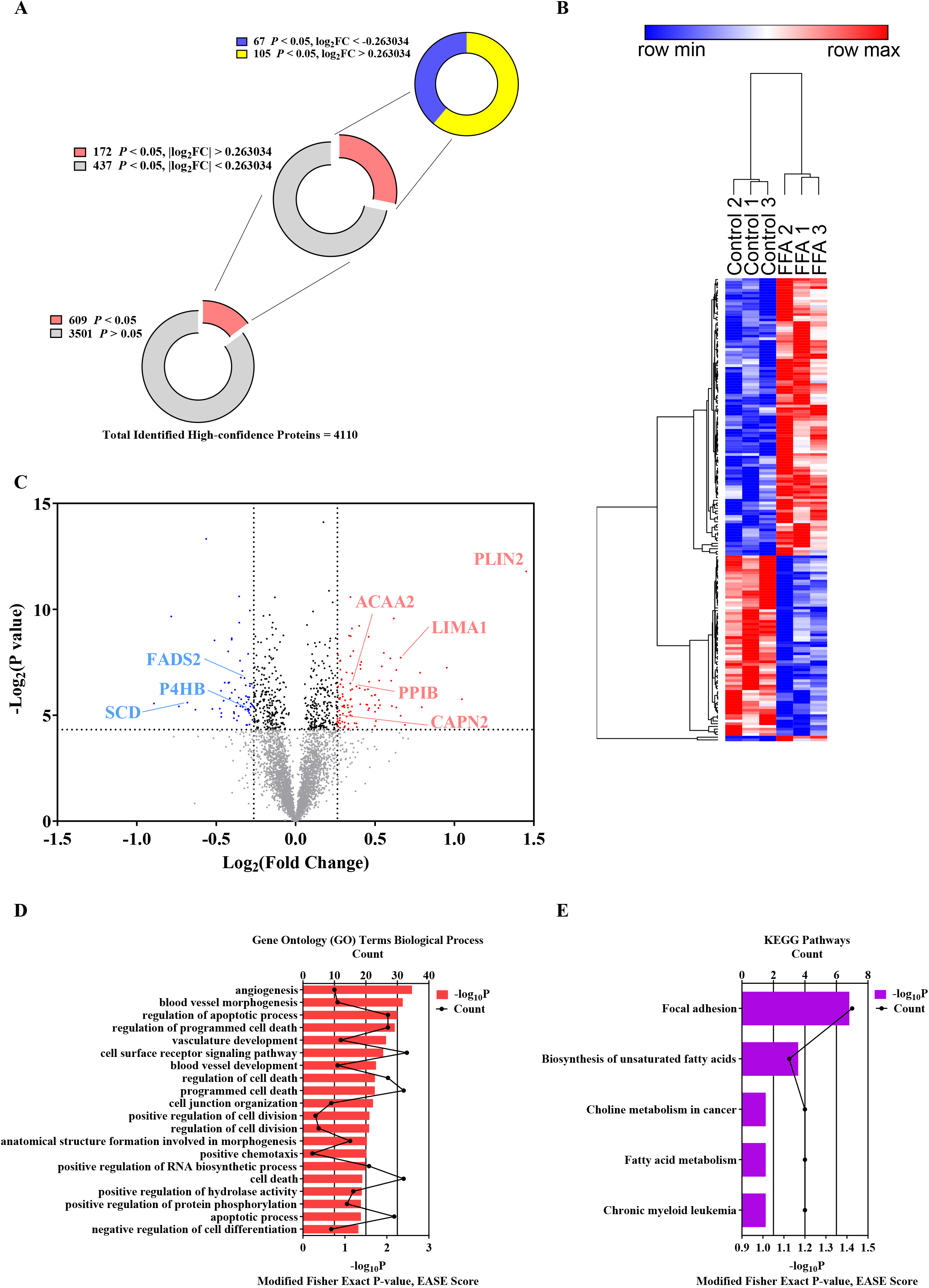
Overview of bioinformatics analysis of FFA-treated Huh7 cells proteomics data. (A) Donut chart of proteomics analysis. A total of 4,110 high-confidence proteins were quantitated by nanoLC-MS/MS in control and FFA treated Huh7 cells. A *p* < 0.05 and |log_2_Fold change| > 0.263034 were selected as the cutoff criteria to identify differentially expressed proteins. 609 proteins showed statistically significant (*p* < 0.05) differential expression between groups. (B) Hierarchical clustering and heatmap of the 172 differentially expressed proteins. Red indicates up-regulation and blue down-regulation. (C) Volcano plot of the proteomics data. The dotted lines show the cutoff limits for *p* and fold-change used. Differentially expressed proteins are highlighted. Red dots represent upregulated proteins and blue dots down-regulated proteins. The dots for differentially regulated proteins for which the transcripts also were differentially regulated are labeled. (D-E) Bar and line charts showing *p* (bars) and count (lines) of top 20 significantly enriched GO BP terms and the significant KEGG pathways from DAVID analysis (*p* < 0.05; modified Fisher’s exact test).

Using bioinformatic analysis, we found significant enrichments (modified Fisher Exact *p* ≤ 0.1) in metabolism process-related GO BP terms and KEGG pathways in FFA-treated cells (Fig. 5D-E). BP included a number of terms related to the apoptotic process, transcription, phosphorylation, cell nucleus, and RNA processing, suggesting effects on cell stress regulation and gene transcription. In accordance with FFA treatment, KEGG pathway analysis revealed enrichment of fatty acid-related pathways (biosynthesis of unsaturated fatty acids and fatty acid metabolism) and choline metabolism. These results are consistent with our transcriptome analysis, which indicated changes in transcript levels for metabolism-related processes in this FFA-induced MAFLD cell model.

By DAVID enrichment analysis, we identified 78 enriched GO BP terms (EASE score < 0.1) in our proteomics analysis. A reduced list of these terms (EASE score <0.1) generated using REVIGO is presented as a graph in Supplementary Figure S1. The analysis of the proteomics enriched BP terms produced a more complex picture (Supplementary Figure S1) than the one obtained for transcriptomics. There are groups related to regulation, differentiation and development, and cell death, respectively. ‘Organelle organization’, ‘cell surface receptor signaling pathway’, and ‘immune system process’ are each linked to all these three groups.

### Comparison of transcriptome and proteomics analyses

Comparison of all the transcripts and the proteins showed little overlap (Fig. 6A). Most of the significant changes were only in either RNA-Seq or proteomics, and only 8 genes were significantly regulated in both. Only one of these (ACAA2) is a mitochondrial protein listed in MitoCarta3.0. Correlation analysis shows a significant but very weak correlation (*p* < 0.0001, Pearson r = 0.084) of the fold changes of transcriptome and proteomics (Fig. 6B). Plotting of the fold changes of the transcripts/proteins quantitated with both proteomics and RNA-Seq methods is shown in Fig. 6C. Four of the genes significantly regulated at transcript and protein level are consistently upregulated, and three are consistently down-regulated; one is up at protein level and down-regulated at the transcript level. The latter (*PPIB)* encodes an ER-localized proline isomerase that catalyzes the cis-trans isomerization of proline residues and may therefore assist protein folding.

**Figure 6:**
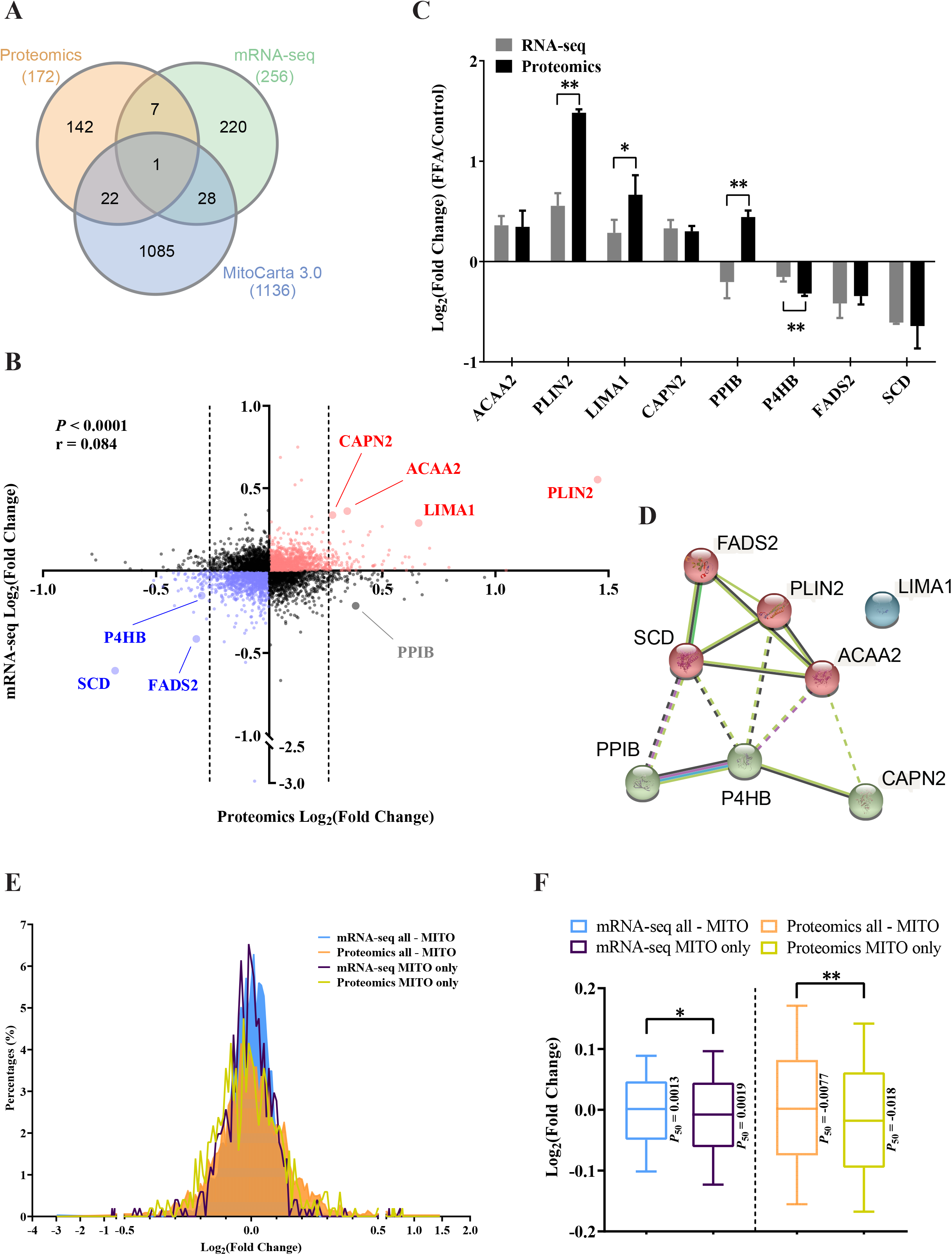
Comparison of regulated transcripts and proteins. (A) Venn diagram showing the overlap between proteomics and transcriptomics and fractions of MitoCarta 3.0 proteins. Fold change plot, red, blue, and grey color dots with labels shows the 8 gene transcripts/proteins fulfilling the criteria of significantly regulated in both proteomics (*p* < 0.05 and fc +/− 20%) and transcriptomics (DESEQ *p* < 0.05). Black stippled lines indicate the threshold cutoffs for proteomics. TPM ≥ 1 was used as a threshold for mRNA-Seq. The statistically significant *p-*value and weakly positive r-value of Pearson’s correlation for genes regulated both at the transcript and protein levels are shown. (C) Fold change graph and (D) protein-protein interaction (PPI) clustered network of the 8 genes/proteins regulated both at protein and transcript level. Network nodes represent proteins, and the colors represent 3 different clusters. Edges represent protein-protein associations, and line color indicates different types of interaction evidence. \**p* < 0.05, \*\**p* < 0.01. (E) Pivot chart showing the frequency of log_2_(fold change) for mRNA-seq and proteomics. TPM ≥ 1 was used as a threshold for mRNA-Seq. (F) Box plot showing the log_2_(fold change) and quartile distributions of genes regulated both at the transcript and protein level. TPM ≥ 1 was used as a threshold for mRNA-Seq. Values are expressed as 10th percentile, 25th percentile, median, 75th percentile, and 90th percentile. \**p* < 0.05, \*\**p* < 0.01.

Remarkably, six of the co-regulated transcripts and proteins are encoding or represent proteins involved in lipid metabolism-related processes: two in fatty acid catabolism (*ACAA2, PLIN2),* two in lipid transport (*LIMA1, P4HB*), and two in the synthesis of unsaturated fatty acids (*FADS2, SCD*). The first two are upregulated and the latter four down-regulated. For a detailed discussion of the functions of these genes, see discussion. In PPI analysis of protein interactions, *FADS2*, *SCD*, *PLIN2,* and *ACAA2* were grouped as a cluster, while *PPIB*, *P4HB*, *CAPN2* were grouped as another cluster. *LIMA1* was grouped as an independent cluster (Fig. 6D).

To investigate whether there was a general regulation of mitochondrial proteins and their coding transcripts, we compared the fold change distributions of all mitochondrial proteins to the fold change distributions of all non-mitochondrial proteins and the fold changes of transcripts encoding mitochondrial proteins to the fold-changes of transcripts encoding non-mitochondrial proteins. The pivot chart shows a marked shift to lower log_2_(fold change) of mitochondrial proteins compared to non-mitochondrial proteins. The log_2_(fold change) distribution of transcripts encoding mitochondrial proteins showed the same trend compared to transcripts encoding non-mitochondrial proteins. (Fig. 6E). We further performed a statistical analysis (Mann Whitney U Test) on the log_2_ (fold change) data between the groups. The log_2_(fold change) of both mitochondrial transcripts compared to transcripts not encoding mitochondrial proteins (*p* < 0.05) and the log_2_(fold change) of mitochondrial proteins compared to non-mitochondrial proteins (*p* < 0.01) were significantly lower. (Fig. 6F). These data indicate that both mitochondrial proteins and transcripts of the MAFLD model cells were expressed at decreased levels in the MAFLD model cells.

## DISCUSSION

In spite of numerous studies, the molecular pathological mechanisms and the order of events in FFA-induced MAFLD are still only insufficiently elucidated, which is why it remains difficult to devise and develop targeted therapies that can delay or counteract the pathogenesis. Deficient handling of the fatty acid surfeit by lipid storage and metabolization systems has in multiple studies been reported to result in mitochondrial dysfunction, altered ROS signaling and damage, thus provoking inflammatory responses and triggering fibrotic signaling(Xiao *et al*, 2017; Luangmonkong *et al*, 2018; Sunny *et al*, 2017). We hypothesized that the FFA-induced MAFLD cell model is an appropriate *in vitro* system to study mitochondrial dysfunction and the underlying gene expression changes thoroughly in order to characterize this system for future drug screening purposes. To this end, we explored the effects of high FFA load by performing a comprehensive analysis of lipid accumulation, mitochondrial phenotypes, cellular energy metabolism, and global gene expression at both transcript and protein levels.

As expected from a previous study using this MAFLD model, treatment with 1200 μM of a 1:2 mixture of the fatty acids palmitate and oleate induced substantial intracellular lipid deposition, which is a remarkable feature of hepatocyte steatosis, and it also appreciably decreased cell viability.

In general, when the extracellular environment changes, cells change their metabolic pattern and thus adapt to the new environment. A high fat load may be balanced by increased mitochondrial fatty acid degradation, producing acetyl-CoA that fuels the tricarboxylic acid cycle. This in turn, increases the flow of reducing equivalents that drive the electron transport chain and oxidative phosphorylation. Using real-time metabolic analysis with the Seahorse analyzer, we explored the effects of 1200 μM FFA-treatment on cellular and mitochondrial energy metabolism. The MAFLD model cells displayed changes in key parameters of mitochondrial and glycolytic ATP production. Compared to control cells, mitochondrial ATP production was significantly increased, and glycolytic ATP production concomitantly decreased. Basal respiration was moderately but significantly increased, while maximal respiration did not show significant changes between control and MAFLD model cells. At the same time, glycolytic fermentation resulting in lactate production was decreased. The increase in respiratory and decrease in glycolytic ATP production led to an unchanged total cellular ATP production in MAFLD model cells. This suggests that the cells maintain an appropriate level of total ATP production. A previous study showed that treatment with a lower dose (750 μM) of a 2:1 mixture of oleic/palmitic acid mixture of rat liver cancer FaO cells for 3 h significantly increased the number of intracellular lipid droplets, but did not significantly alter basal respiration, maximal respiration, or ATP production (Vecchione *et al*, 2017). The inconsistent results for basal respiration may be due to different concentrations of FFAs or, more likely, the different incubation times with the FFAs: 24 hours in our experiments and 3 hours in the other study.

Oxidative stress has been strongly indicated to play an important role in MAFLD (Svegliati-Baroni *et al*, 2019; Mansouri *et al*, 2018), and antioxidants have been suggested as one therapeutic option for MAFLD treatment. The leakage of electrons from the mitochondrial electron transport chain that react with oxygen to form superoxide is a major source of oxidative stress (Mercado-Uribe *et al*, 2020; Majumder *et al*, 2021). The increased flow of electrons through the respiratory chain may thus be expected to increase the production of mitochondrial superoxide. As we only observed a moderate increase in mitochondrial respiration, we also did not expect a strong increase in mitochondrial superoxide levels. Indeed, our high-throughput imaging flow cytometry measurements showed no significant changes in mitochondrial superoxide levels both as per cell or normalized to mitochondrial volume. This indicates that either the production of superoxide is not increased or that higher superoxide production is balanced by increased activity of superoxide dismutase 2 (SOD2). The latter would both scavenge damage to macromolecules and increase redox signaling, e.g. via the KEAP1-NRF2 system, a redox-sensitive signaling system mediating the response to oxidant stress (Suzuki *et al*, 2019). Previous studies have suggested that the levels of oxidative stress in the liver vary in different MAFLD stages. Interestingly, while many studies described increased ROS in MAFLD (Upadhyay *et al*, 2020; Mansouri *et al*, 2018), it has also been discussed that there may be decreased ROS levels at specific stages of MAFLD (Piccinin *et al*, 2019). Our study indicates that increased superoxide levels inside mitochondria are not an important factor in the early stages of MAFLD in hepatocytes.

Given that the high fat load only caused a moderate shift in mitochondrial and glycolytic ATP production without changing the rate of total ATP production and the finding that mitochondrial superoxide levels remained unchanged, one may expect that the 24 hours of high fat exposure may have triggered changes in gene expression. Transcriptome analysis revealed a moderate number, 256, of significant transcript level changes. This prominently included genes associated with lipid metabolism-related terms, but also signaling pathways like PPAR and AKT signaling. Network analysis of the regulated transcripts revealed lipid handling and responses and signaling as common denominators and connected via the central term ‘metabolic process’. This suggests that the major primary effects of high fat treatment impact the levels of genes involved in metabolism. Finally, GSEA fine analysis of the mRNA-Seq data showed significant enrichment of the chemokine receptor binding gene set, which is an evidence for initiation of the inflammation.

Proteome analysis also showed a fair number of differentially regulated proteins, 172, in the MAFLD cell model. However, the connection to lipid metabolism was less clear. Interestingly, we found a general down-regulation of mitochondrial proteins in FFA treated cells suggesting less mitochondria content. This was supported by comparing the transcripts encoding mitochondrial proteins with those encoding non-mitochondrial proteins. However, here the difference was less pronounced. Using MitoTracker Green in the HTIFC experiment, we found that the mitochondrial volume of FFA treated cells did not change significantly. A possible explanation might be an accelerated turnover of mitochondrial proteins by mitochondrial proteases without affecting mitochondrial volume. The further analysis of the multi-omics data through the mitochondrial gene set MitoPathways 3.0 in MitoCarta 3.0 showed the expression levels of genes and proteins within the mitophagy, mitochondrial fission, mitochondrial fusion, and tricarboxylic acid cycle data sets were not altered (data not shown). However, interestingly, 10.8% of the protein or transcript expression levels changed significantly in the mitochondrial respiratory chain set (data not shown). Taken together, this may indicate that a specific subset of the mitochondrial proteome is downregulated. In this regard, further research is needed.

Clustering analysis of the differentially regulated proteins into networks revealed hubs of proteins related to regulation, differentiation and development, and cell death. The clusters are highly interconnected with each other. When comparing the significantly regulated transcripts with the significantly regulated proteins, there was only very little overlap. There is a weak correlation between all transcripts/proteins quantitated in both analyses. Transcript and protein levels correlate typically not very well as transcript and protein turnover, translation efficiency, and other parameters affect the levels of transcripts and proteins differently (Edfors *et al*, 2016; Moritz *et al*, 2019; Bauernfeind & Babbitt, 2017).

When focusing on the genes consistently regulated both at the transcript and protein level, it is remarkable that six of the seven genes have a link to lipid metabolism. ACAA2 is a thiolase that catalyzes the last step in the mitochondrial catabolism of medium-chain fatty acids, producing acetyl-CoA. In the reverse reaction, it catalyzes the first step in the ketone body and cholesterol synthesis pathways. We find ACAA2 gene expression upregulated; an increase in fatty acid oxidation capacity could feed the TCA cycle and the respiratory chain explaining the increased mitochondrial ATP production rate that we observed in the Seahorse metabolic measurements.

Perilipin 2 (PLIN2), also upregulated, is the only constitutive and ubiquitously expressed lipid droplet protein of the perilipin family and it represents a marker for the number and volume of lipid droplets (Tsai *et al*, 2017). Plin2^-/-^ mice have an approximately 60% reduction in triglyceride content. PLIN2 deletion in the adrenal cortex has been shown to increase cholesterol content in lipid droplets (Tsai *et al*, 2017). Furthermore, PLIN2 downregulation stimulates triglyceride catabolism via autophagy while PLIN2 overexpression protects against autophagy. The enhanced autophagy in *Plin2*^-/-^ mice protects against severe ER stress-induced hepatosteatosis and hepatocyte apoptosis (Tsai *et al*, 2017). PLIN2 up-regulation, as we observed in the FFA treated cells, may be a response to increase the storage capacity for fatty acids in lipid droplets. The beneficial effect on fatty acid storage capacity may, however, come at a price that triggers detrimental responses.

Notably, *SCD* and *FADS2,* both encoding fatty acid desaturases that introduce double bonds into fatty acids are downregulated. SCD introduces the first double bond into saturated fatty acids generating monounsaturated fatty acids, whereas FADS2 in turn catalyzes the further desaturation of monounsaturated fatty acid precursors (Koletzko *et al*, 2019). Mono- and polyunsaturated fatty acids (MUFAs and PUFA’s, respectively) contribute to cell growth, survival, differentiation, metabolic regulation, and signal transduction. Overexpression of SCD is implicated in metabolic diseases such as diabetes and non-alcoholic fatty liver disease (Kikuchi & Tsukamoto, 2020). SCD facilitates metabolic reprogramming in cancer mediated, at least in part, by regulation of AKT, AMPK, and NF-kB via MUFAs. Interesting with respect to our Huh7 MALFD model, treatment of Huh7 cells with the mTOR inhibitors Torin 1 or rapamycin decreased expression of FADS2 and consequently reduced new synthesis of the desaturated fatty acids palmitoleate, palmitate, and sapienate (Triki *et al*, 2020). 2/3 of our FFA treatment consists of oleic acid, a monounsaturated fatty acid. Down-regulation of SCD, which generates oleic acid from stearic acid, makes thus sense. The observation that FADS2 that catalyzes further desaturation also is down-regulated suggests that expression of the two genes is co-regulated. As both desaturases are down-regulated, this may indicate a response that limits the synthesis of unsaturated fatty acids.

The upregulated *LIMA1* gene encodes an actin binding protein involved in regulation and dynamics of the actin cytoskeleton. It has also a connection to lipid metabolism as it is implicated in recycling of NPLC1L1, a receptor protein responsible for cholesterol uptake from the intestinal lumen and liver canicular space and thus contributes to cholesterol homeostasis (Luo *et al*, 2020). It increases the number and size of actin stress fibers and inhibits membrane ruffling, a characteristic feature of migrating cells (Maul *et al*, 2003).

The *P4HB* gene encodes a multifunctional protein that catalyzes the formation, breakage and rearrangement of disulfide bonds. At the cell surface, it seems to act as a reductase that cleaves disulfide bonds of proteins attached to the cell and may, in this way, cause structural modifications of exofacial proteins. Inside the cell, P4HB seems to form/rearrange disulfide bonds of nascent proteins. Notable in the context of MAFLD, P4HB also acts as a structural subunit of various enzymes, one of which is microsomal triacylglycerol transfer protein (MTTP). MTTP catalyzes the transport of triglyceride, cholesteryl ester, and phospholipid between phospholipid surfaces (Lazaris *et al*, 2021).

Taken together, the six ‘tip of the iceberg’ regulated genes encoding lipid metabolism, transport, and handling related proteins suggests that lipid homeostasis is strongly disturbed in the MAFLD model resulting in compensatory but also potentially damaging changes when the disturbance of the lipid homeostasis is persistent as seen in MAFLD patients.

Finally, *CAPN2*, which we find upregulated, has no obvious relationship to lipid metabolism. Like *LIMA1,* it encodes a protein involved in actin cytoskeleton dynamics. When localized in the cytosol, it catalyzes limited proteolysis of substrates involved in cytoskeletal remodeling and signal transduction. Upon Ca2^+^ binding, CAPN2 translocates to the plasma membrane and is involved in the degradation of the extracellular matrix. It has been found that its up-regulation upon HBV infection induces the expression of hepatic fibrosis markers (Feng *et al*, 2020). Its upregulation in our MAFLD model may indicate the beginning induction of fibrosis, which occurs in the transitions from the fatty liver via NASH to fibrosis.

## Conclusions

In this study, we investigated the characteristics of cellular energy metabolism adaptations and gene expression reprogramming in a 1200 μM FFA-induced MAFLD model. We found that this cell model is an appropriate *in vitro* model to study the role of mitochondria, fatty acid metabolism, glycolysis, and multiple metabolic pathways in MAFLD. This model can also be used for drug screening (including traditional Chinese medicine) and research for treatment mechanisms for fatty acid metabolism disorders, mitochondrial dysfunction, and abnormal glycolysis.

Although *in vitro* models cannot replace animal models, *in vitro* models have economic and time advantages in screening multiple drugs at the same time. Furthermore, because one can test one cell type at a time, it excludes the influence of other cell types in the liver, and the effective mechanisms for MAFLD drugs can be analyzed specifically for that cell type.

## List of abbreviations

BSA: bovine serum albumin
BP: biological process
CC: cellular component
DMEM: Dulbecco’s Modified Eagle Medium
FBS: fetal bovine serum
FFA: free fatty acid
GO: gene ontology
GSEA: gene set enrichment analysis
HCC: hepatocellular carcinoma
HTIFC: high-throughput imaging flow cytometry
KEGG: Kyoto encyclopedia of genes and genomes
LC-MS/MS: liquid chromatography tandem mass spectrometry
MAFLD: metabolic associated fatty liver diseases
MF: molecular function
NAFLD: non-alcoholic fatty liver disease
OCR: oxygen consumption rate
OXPHOS: oxidative phosphorylation
PER: proton efflux rate
PPI: protein-protein interaction
PSM: peptide spectrum matches
ROS: reactive oxygen species
Rot/AA: rotenone plus antimycin A
SCX: strong cation exchange
TMM: trimmed mean of M value
TPM: transcripts per million

## Supplementary material

**Supplementary Figure S1.**
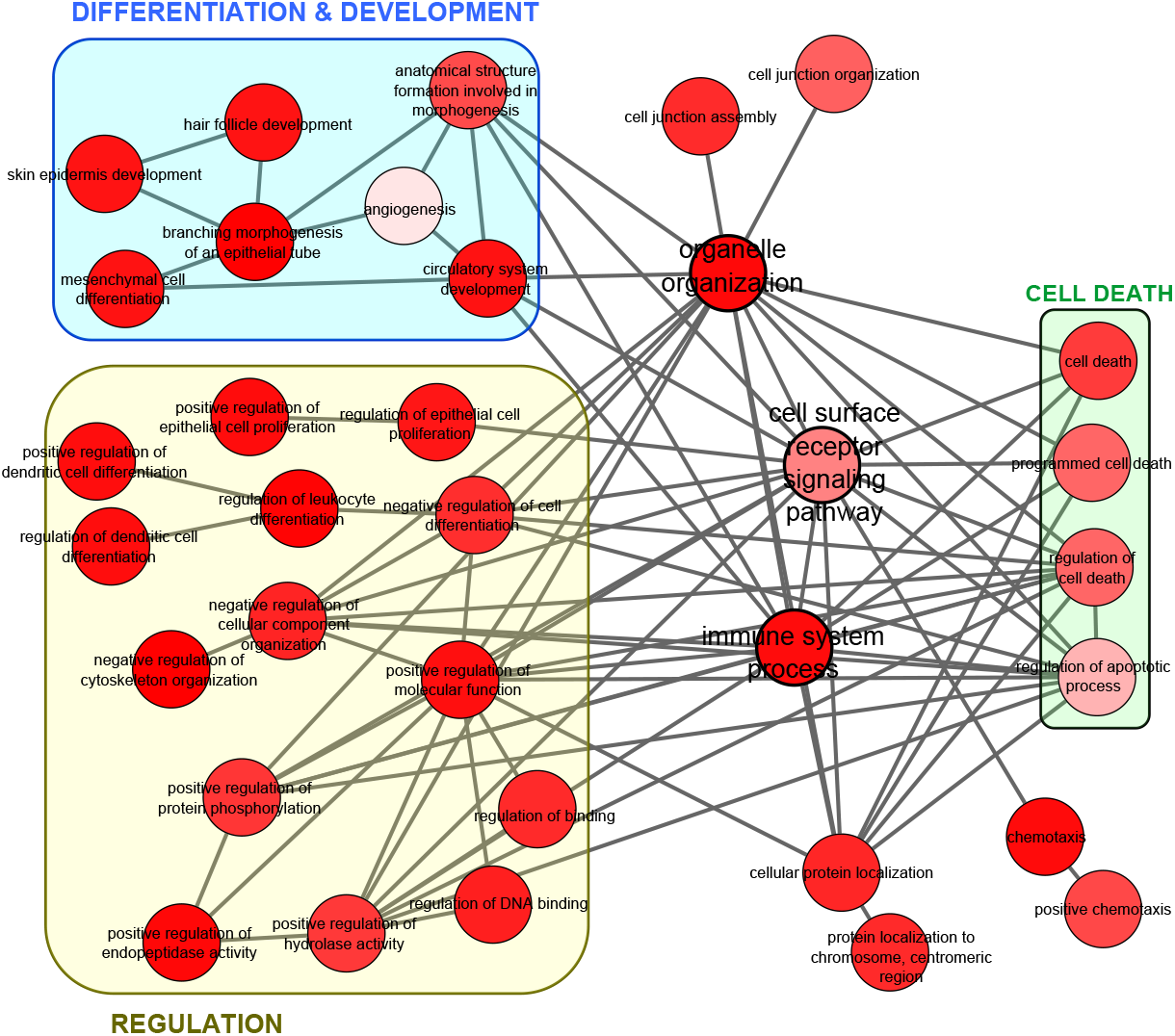
Enriched GO BP terms in proteomics differentially expressed genes displayed after elimination of redundant GO terms using REVIGO (Shannon *et al*, 2003). Settings: GO BP terms with EASE score (modified Fisher’s exact Test; *p* < 0.1) and the associated *p-*value were used as input for REVIGO. The REVIGO analysis was performed with similarity = small (0.5), species = homo sapiens, semantic similarity measure = Jiang and Conrath. The resulting interactive graph was edited using Cytoscape. Term groups related to ‘regulation’ and ‘differentiation & development’ are boxed. The three terms ‘organelle organization’, ‘cell surface receptor signaling pathway’, and ‘immune system process’ are highlighted by different colors of the bubble border, and the lines of their immediate connections are displayed in the border color.

## ACKNOWLEDGMENTS

This work was supported by grants from National Natural Science Foundation of China (No. 81974558) and International Program for Postgraduates, Guangzhou University of Chinese Medicine.

## REFERENCES

Anders S & Huber W (2010) Differential expression analysis for sequence count data. Genome Biology 11: R106

Anstee QM, Reeves HL, Kotsiliti E, Govaere O & Heikenwalder M (2019) From NASH to HCC: current concepts and future challenges. Nat Rev Gastroenterol Hepatol 16: 411–428

Bauernfeind AL & Babbitt CC (2017) The predictive nature of transcript expression levels on protein expression in adult human brain. BMC Genomics 18: 1–11

Chavez-Tapia NC, Rosso N & Tiribelli C (2012) Effect of intracellular lipid accumulation in a new model of non-alcoholic fatty liver disease. BMC Gastroenterology 12: 20

Chen Z, Tian R, She Z, Cai J & Li H (2020) Role of oxidative stress in the pathogenesis of nonalcoholic fatty liver disease. Free Radical Biology and Medicine 152: 116–141

Cheng D, Yang A, Thomas H & Monjardino J (1993) Characterization of stable hepatitis delta expressing hepatoma cell lines: effect of HDAg on cell growth. Progress in clinical and biological research 382: 149–153

Cholankeril G, Perumpail RB, Pham EA, Ahmed A & Harrison SA (2016) Nonalcoholic Fatty Liver Disease: Epidemiology, Natural History, and Diagnostic Challenges. Hepatology 64: 954

Crowley LC, Marfell BJ & Waterhouse NJ (2016) Analyzing cell death by nuclear staining with Hoechst 33342. Cold Spring Harbor Protocols 2016: 778–781

Edfors F, Danielsson F, Hallström BM, Käll L, Lundberg E, Pontén F, Forsström B & Uhlén M (2016) Gene‐specific correlation of RNA and protein levels in human cells and tissues. Molecular Systems Biology 12: 883

Eslam M, Sanyal AJ, George J, Sanyal A, Neuschwander-Tetri B, Tiribelli C, Kleiner DE, Brunt E, Bugianesi E, Yki-Järvinen H, et al (2020) MAFLD: A Consensus-Driven Proposed Nomenclature for Metabolic Associated Fatty Liver Disease. Gastroenterology 158: 1999–2014.e1

Feng R, Du W, Lui P, Zhang J & Liu Y (2020) CAPN2 acts as an indicator of hepatitis B virus to induce hepatic fibrosis. Journal of Cellular Biochemistry 121: 2428–2436

Gariani K, Ryu D, Menzies KJ, Yi HS, Stein S, Zhang H, Perino A, Lemos V, Katsyuba E, Jha P, et al (2017) Inhibiting poly ADP-ribosylation increases fatty acid oxidation and protects against fatty liver disease. Journal of Hepatology 66: 132–141

Hardy T, Oakley F, Anstee QM & Day CP (2016) Nonalcoholic Fatty Liver Disease: Pathogenesis and Disease Spectrum. Annual Review of Pathology: Mechanisms of Disease 11: 451–496

Heberle H, Meirelles VG, da Silva FR, Telles GP & Minghim R (2015) InteractiVenn: A web-based tool for the analysis of sets through Venn diagrams. BMC Bioinformatics 16: 169

Huang DW, Sherman BT & Lempicki RA (2009a) Systematic and integrative analysis of large gene lists using DAVID bioinformatics resources. Nature Protocols 4: 44–57

Huang DW, Sherman BT & Lempicki RA (2009b) Bioinformatics enrichment tools: Paths toward the comprehensive functional analysis of large gene lists. Nucleic Acids Research 37: 1–13

Kikuchi K & Tsukamoto H (2020) Stearoyl-CoA desaturase and tumorigenesis. Chemico-Biological Interactions 316 doi:10.1016/j.cbi.2019.108917 [PREPRINT]

Koletzko B, Reischl E, Tanjung C, Gonzalez-Casanova I, Ramakrishnan U, Meldrum S, Simmer K, Heinrich J & Demmelmair H (2019) FADS1 and FADS2 Polymorphisms Modulate Fatty Acid Metabolism and Dietary Impact on Health. Annual review of nutrition 39: 21–44 doi:10.1146/annurev-nutr-082018-124250 [PREPRINT]

Lazaris V, Hatziri A, Symeonidis A & Kypreos KE (2021) The Lipoprotein Transport System in the Pathogenesis of Multiple Myeloma: Advances and Challenges. Frontiers in Oncology 11 doi:10.3389/fonc.2021.638288 [PREPRINT]

Levin Y (2011) The role of statistical power analysis in quantitative proteomics. Proteomics 11: 2565–2567

Li M, Xu C, Shi J, Ding J, Wan X, Chen D, Gao J, Li C, Zhang J, Lin Y, et al (2017) Fatty acids promote fatty liver disease via the dysregulation of 3-mercaptopyruvate sulfurtransferase/hydrogen sulfide pathway. Gut 67

Liberzon A, Birger C, Thorvaldsdóttir H, Ghandi M, Mesirov JP & Tamayo P (2015) The Molecular Signatures Database Hallmark Gene Set Collection. Cell Systems 1: 417–425

Liberzon A, Subramanian A, Pinchback R, Thorvaldsdóttir H, Tamayo P & Mesirov JP (2011) Molecular signatures database (MSigDB) 3.0. Bioinformatics 27: 1739–1740

Luangmonkong T, Suriguga S, Mutsaers HAM, Groothuis GMM, Olinga P & Boersema M (2018) Targeting Oxidative Stress for the Treatment of Liver Fibrosis. Rev Physiol Biochem Pharmacol 175: 71–102

Luo J, Yang H & Song BL (2020) Mechanisms and regulation of cholesterol homeostasis. Nature Reviews Molecular Cell Biology 21: 225–245 doi:10.1038/s41580-019-0190-7 [PREPRINT]

Majumder D, Nath P, Debnath R & Maiti D (2021) Understanding the complicated relationship between antioxidants and carcinogenesis. Journal of Biochemical and Molecular Toxicology 35 doi:10.1002/jbt.22643 [PREPRINT]

Mansouri A, Gattolliat CH & Asselah T (2018) Mitochondrial Dysfunction and Signaling in Chronic Liver Diseases. Gastroenterology 155: 629–647 doi:10.1053/j.gastro.2018.06.083 [PREPRINT]

Maul RS, Song Y, Amann KJ, Gerbin SC, Pollard TD & Chang DD (2003) EPLIN regulates actin dynamics by cross-linking and stabilizing filaments. Journal of Cell Biology 160: 399–407

Mercado-Uribe H, Andrade-Medina M, Espinoza-Rodríguez JH, Carrillo-Tripp M & Scheckhuber CQ (2020) Analyzing structural alterations of mitochondrial intermembrane space superoxide scavengers cytochrome-c and SOD1 after methylglyoxal treatment. PLoS ONE 15

von Mering C, Huynen M, Jaeggi D, Schmidt S, Bork P & Snel B (2003) STRING: A database of predicted functional associations between proteins. Nucleic Acids Research 31: 258–261 doi:10.1093/nar/gkg034 [PREPRINT]

Mootha VK, Lindgren CM, Eriksson KF, Subramanian A, Sihag S, Lehar J, Puigserver P, Carlsson E, Ridderstråle M, Laurila E, et al (2003) PGC-1α-responsive genes involved in oxidative phosphorylation are coordinately downregulated in human diabetes. Nature Genetics 34: 267–273

Moritz CP, Mühlhaus T, Tenzer S, Schulenborg T & Friauf E (2019) Poor transcript-protein correlation in the brain: negatively correlating gene products reveal neuronal polarity as a potential cause. Journal of Neurochemistry 149: 582–604

Palmfeldt J, Vang S, Stenbroen V, Pedersen CB, Christensen JH, Bross P & Gregersen N (2009) Mitochondrial proteomics on human fibroblasts for identification of metabolic imbalance and cellular stress. Proteome Science 7

Paternoster V, Svanborg M, Edhager AV, Rajkumar AP, Eickhardt EA, Pallesen J, Grove J, Qvist P, Fryland T, Wegener G, et al (2019) Brain proteome changes in female Brd1+/− mice unmask dendritic spine pathology and show enrichment for schizophrenia risk. Neurobiology of Disease 124: 479–488

Patterson J, Carpenter EJ, Zhu Z, An D, Liang X, Geng C, Drmanac R & Wong GKS (2019) Impact of sequencing depth and technology on de novo RNA-Seq assembly. BMC Genomics 20

Piccinin E, Villani G & Moschetta A (2019) Metabolic aspects in NAFLD, NASH and hepatocellular carcinoma: the role of PGC1 coactivators. Nature Reviews Gastroenterology and Hepatology 16: 160–174 doi:10.1038/s41575-018-0089-3 [PREPRINT]

Rao J, Peng L, Liang X, Jiang H, Geng C, Zhao X, Liu X, Fan G, Chen F & Mu F (2020) Performance of copy number variants detection based on whole-genome sequencing by DNBSEQ platforms. BMC Bioinformatics 21

Rath S, Sharma R, Gupta R, Ast T, Chan C, Durham TJ, Goodman RP, Grabarek Z, Haas ME, Hung WHW, et al (2021) MitoCarta3.0: An updated mitochondrial proteome now with sub-organelle localization and pathway annotations. Nucleic Acids Research 49: D1541–D1547

Reimand J, Isserlin R, Voisin V, Kucera M, Tannus-Lopes C, Rostamianfar A, Wadi L, Meyer M, Wong J, Xu C, et al (2019) Pathway enrichment analysis and visualization of omics data using g:Profiler, GSEA, Cytoscape and EnrichmentMap. Nature Protocols 14: 482–517

Reynolds JE, Li J & Eastman A (1996) Detection of apoptosis by flow cytometry of cells simultaneously stained for intracellular pH (carboxy SNARF-1) and membrane permeability (Hoechst 33342). Cytometry 25: 349–357

Robinson MD & Oshlack A (2010) A scaling normalization method for differential expression analysis of RNA-seq data. Genome Biology 11: R25

Ruhanen H, Nidhina Haridas PA, Eskelinen EL, Eriksson O, Olkkonen VM & Käkelä R (2017) Depletion of TM6SF2 disturbs membrane lipid composition and dynamics in HuH7 hepatoma cells. Biochimica et Biophysica Acta - Molecular and Cell Biology of Lipids 1862: 676–685

Santhekadur PK, Kumar DP & Sanyal AJ (2018) Preclinical models of non-alcoholic fatty liver disease. Journal of Hepatology 68: 230–237 doi:10.1016/j.jhep.2017.10.031 [PREPRINT]

Shannon P, Markiel A, Ozier O, Baliga NS, Wang JT, Ramage D, Amin N, Schwikowski B & Ideker T (2003) Cytoscape: A software Environment for integrated models of biomolecular interaction networks. Genome Research 13: 2498–2504

Su G, Morris JH, Demchak B & Bader GD (2014) Biological Network Exploration with Cytoscape 3. Current Protocols in Bioinformatics 2014: 8.13.1–8.13.24

Subramanian A, Tamayo P, Mootha VK, Mukherjee S, Ebert BL, Gillette MA, Paulovich A, Pomeroy SL, Golub TR, Lander ES, et al (2005) Gene set enrichment analysis: A knowledge-based approach for interpreting genome-wide expression profiles. Proceedings of the National Academy of Sciences of the United States of America 102: 15545–15550

Sunny NE, Bril F & Cusi K (2017) Mitochondrial Adaptation in Nonalcoholic Fatty Liver Disease: Novel Mechanisms and Treatment Strategies. Trends Endocrinol Metab 28: 250–260

Supek F, Bošnjak M, Škunca N & Šmuc T (2011) Revigo summarizes and visualizes long lists of gene ontology terms. PLoS ONE 6: e21800

Suzuki T, Muramatsu A, Saito R, Iso T, Shibata T, Kuwata K, Kawaguchi S-I, Iwawaki T, Adachi S, Suda H, et al (2019) Molecular Mechanism of Cellular Oxidative Stress Sensing by Keap1. Cell reports 28: 746–758.e4

Svegliati-Baroni G, Pierantonelli I, Torquato P, Marinelli R, Ferreri C, Chatgilialoglu C, Bartolini D & Galli F (2019) Lipidomic biomarkers and mechanisms of lipotoxicity in non-alcoholic fatty liver disease. Free Radical Biology and Medicine 144: 293–309 doi:10.1016/j.freeradbiomed.2019.05.029 [PREPRINT]

Szklarczyk D, Gable AL, Lyon D, Junge A, Wyder S, Huerta-Cepas J, Simonovic M, Doncheva NT, Morris JH, Bork P, et al (2019) STRING v11: Protein-protein association networks with increased coverage, supporting functional discovery in genome-wide experimental datasets. Nucleic Acids Research 47: D607–D613

Triki M, Rinaldi G, Planque M, Broekaert D, Winkelkotte AM, Maier CR, Janaki Raman S, Vandekeere A, Van Elsen J, Orth MF, et al (2020) mTOR Signaling and SREBP Activity Increase FADS2 Expression and Can Activate Sapienate Biosynthesis. Cell Reports 31

Tsai TH, Chen E, Li L, Saha P, Lee HJ, Huang LS, Shelness GS, Chan L & Chang BHJ (2017) The constitutive lipid droplet protein PLIN2 regulates autophagy in liver. Autophagy 13: 1130–1144

Upadhyay KK, Jadeja RN, Vyas HS, Pandya B, Joshi A, Vohra A, Thounaojam MC, Martin PM, Bartoli M & Devkar R V. (2020) Carbon monoxide releasing molecule-A1 improves nonalcoholic steatohepatitis via Nrf2 activation mediated improvement in oxidative stress and mitochondrial function. Redox Biology 28

Vecchione G, Grasselli E, Cioffi F, Baldini F, Oliveira PJ, Sardão VA, Cortese K, Lanni A, Voci A, Portincasa P, et al (2017) The Nutraceutic Silybin Counteracts Excess Lipid Accumulation and Ongoing Oxidative Stress in an In Vitro Model of Non-Alcoholic Fatty Liver Disease Progression. Frontiers in Nutrition 4

Wagner GP, Kin K & Lynch VJ (2012) Measurement of mRNA abundance using RNA-seq data: RPKM measure is inconsistent among samples. Theory in Biosciences 131: 281–285

Xiao M, Zhong H, Xia L, Tao Y & Yin H (2017) Pathophysiology of mitochondrial lipid oxidation: Role of 4-hydroxynonenal (4-HNE) and other bioactive lipids in mitochondria. Free Radic Biol Med 111: 316–327

Younossi ZM (2019) Non-alcoholic fatty liver disease – A global public health perspective. Journal of Hepatology 70: 531–544

Younossi ZM, Loomba R, Rinella ME, Bugianesi E, Marchesini G, Neuschwander-Tetri BA, Serfaty L, Negro F, Caldwell SH, Ratziu V, et al (2018) Current and future therapeutic regimens for nonalcoholic fatty liver disease and nonalcoholic steatohepatitis. Hepatology 68: 361–371

Yue B, Yang H, Wu J, Wang J, Ru W, Cheng J, Huang Y, Lei C, Lan X & Chen H (2020) Characterization and transcriptome analysis of exosomal and nonexosomal rnas in bovine adipocytes. International Journal of Molecular Sciences 21: 1–17

Zhu X, Feng J, Fu W, Shu X, Wan X & Liu J (2020) Effects of cisplatin on the proliferation, invasion and apoptosis of breast cancer cells following β-catenin silencing. International Journal of Molecular Medicine 45: 1838–1850

